# Impact of gene selection criteria on transcriptomic ontology-based point of departure estimates

**DOI:** 10.1101/2023.10.20.561869

**Authors:** Michael B. Black, Alina Y. Efremenko, A. Rasim Barutcu, Patrick D. McMullen

## Abstract

Apical effects are typically associated with changes in gene expression, which allows for the use of short-term in life transcriptomic studies to derive biologically relevant points of departure (POD). These methods offer cost savings over conventional toxicology assessments and can derive data from very short-term studies where apical effects may not yet be present. When there is limited or insufficient data for a conventional POD assessment, a transcriptomic screen could provide valuable data for deriving a cellular bioactivity POD for chemical screening and hazard assessment. We used existing transcriptomic data from published 5-day rat in vivo kidney and liver exposures to examine the effect of differential gene expression metrics for the selection of genes used for ontology pathway-based POD derivation. Williams Trend Test (WTT) indicate no gene expression dose-response in 6 instances and ANOVA in one, while DESeq2 detected differentially expressed genes in all instances. The three statistical metrics produced consistent POD values. One chemical (PFOA in liver) showed ontology enrichment indicative of a cytotoxic response at the highest dose, emphasizing the effect which too high a dose can have on the derivation of POD values if such response is not accounted for. Whether the choice of a gene selection metric combining both a statistical significance criterion as well as a minimum magnitude of change threshold affects the sensitivity of POD values depends on the specifics of the dose-response. Existing alternative and complementary analyses could be utilized with existing analyses pipelines to better inform analytical decisions when using transcriptomics and BMD for point of departure determinations.

## Introduction

Application of benchmark dose (BMD) analyses to gene expression data from transcriptomic experiments (microarray, qPCR, RNA-Seq, single cell sequence) is emerging as a promising tool in 21^st^ century toxicology (Harrill et al., 2019). BMD analyses of whole transcriptome gene expression data was developed over a decade ago (Thomas et al., 2007; Yang et al., 2007). When combined with BMD analyses and ontology enrichment, transcriptomic data produces POD results that are highly correlated with apical endpoints, even from very short-term exposures (Thomas et al., 2013b). This provides an opportunity to drastically decrease cost and increase throughput of chemical hazard screening by using short term in vivo or in vitro transcriptomic dose-response experiments to derive POD values for assessment.

Transcriptomic data measures changes in gene expression upon exposing cells from in vivo or in vitro dose-response experiments, relative to unexposed control samples. As changes in gene expression are the precursor for functional changes in cell biology, even short-term exposures are useful for detecting changes rapidly for use in assessing the hazard of a chemical (Thomas et al., 2009; Thomas et al., 2013a; Thomas et al., 2013b). Changes in gene expression can be related to cellular functional biology through the use of ontologies, gene set enrichment analyses (GSEA) and other tools to filter lists of genes with altered expression through known functional relationships of genes, their gene products and the role they have in cellular function.

A challenge with transcriptomic data is the inherent variability in levels of mRNA expression in any dataset due to biological variation in expression between samples and even between cells over time in the population of cells sampled from a tissue or in vitro cell population (Jong et al., 2019). In order to determine if a gene’s expression is differentially altered due to exposure to a chemical, significance is determined by a statistical metric to determine the probability of rejecting a null hypothesis. In differential gene expression, the null hypothesis is that changes in expression after chemical exposure are not different than expression levels observed in untreated samples. Significance is usually presented as a false discovery rate corrected p-value or FDR. At the same time genes are assessed for minimum magnitude of change in expression to further ensure confidence in the measured change. Historically this method of combining a statistical and a magnitude of change threshold produces genes lists that are consistent with regard to their representative functional role in a cellular ontology context regardless of differences in the specific genes detected (Guo et al., 2006). Significant genes are used in an over-representation analysis with an ontology to determine which ontology categories share elements with the significant gene list that would not be expected by chance alone (frequently done using a Fisher’s exact test of input gene list relative to the defined gene lists for the ontology categories). In this way, changes in gene expression due to exposure to a chemical can be linked to the biology pathways those genes are associated with. This information in turn can be used for the inference of a cellular mode of action (MOA) (Andersen et al., 2018b; Joseph, 2017; LaRocca et al., 2020a). Alternatives to this method include various forms of Gene Set Enrichment Analyses (GSEA) which do not pre-select list for enrichment but instead compare expression data to pre-defined list of genes with some coordinated underlying biological activity such as shared transcriptional regulation or shared apical outcomes (Chen et al., 2013; Liberzon et al., 2011; Subramanian et al., 2005). A problem with this latter approach is the derivation of appropriate gene lists which may be limiting for many industrial or agricultural chemicals with no intentional human cellular target or which show a diverse cellular response. However, recent analyses indicate that the method may work well, and is particularly amenable to large scale screening studies (Harrill et al., 2021).

A BMD analyses to derive biologically relevant POD values begins with fitting dose-response models to normalized expression data derived for each gene and each sample in a transcriptomic dose-response experiment. Genes are pre-filtered using some metric of differential gene expression to exclude genes not significantly altered as a response to exposure. Multiple BMD models are fit to the individual gene expression data and a best fitting model selected statistically Tools to combine functional ontology enrichment with transcriptomic BMD analyses allow the use of gene-based BMD values to be filtered through cellular functional biology information to then use functional pathway summary BMD values as PODs in chemical hazard screening experiments (Ewald et al., 2020; Phillips et al., 2019; Serra et al., 2020). These approaches are now being increasingly investigated as alternative to conventional toxicology in vivo experiments to derive POD values as transcriptomic experiments can be done for less cost, with higher throughput and shorter exposures than conventional experiments (Baltazar et al., 2020; Gwinn et al., 2020; Johnson et al., 2020; LaRocca et al., 2020b; Martinez et al., 2020; Page-Lariviere et al., 2019; Pouzou et al., 2020; White et al., 2020).

Over the past few years effort has been devoted to developing best practices for the application of transcriptomic BMD analyses for POD derivation (Haber et al., 2018; Jensen et al., 2019; National Toxicology Program, 2018; Toxicogenomics Working Group, 2018). The issues are primarily ones of applying appropriate and rigorous assessments of data at various steps in the analyses to ensure that the results appropriate for the purpose of assessing biological hazard. Not all dose-response experiments will successfully capture an appropriate transcriptomic response for use in BMD analyses. Abrupt inflection in DGE at high dose is not uncommon in transcriptomic screening studies (where dose range finding may not be possible) and may represent a situation of no significant dose-response until a threshold is crossed at high dose. Such threshold response is not represented in most current software for transcriptomic BMD analyses. Threshold response may be well modeled using various bent-hyperbola, spline or broken hockey stick models but if not identified can confound POD derivation due to non-specific gene changes related to cytotoxicity or other high dose stress response not directly relevant to the compounds MOA (Judson et al., 2016; Spassova, 2019).

In their 2018 research report, the National Toxicology program recommended pre-testing each gene expression data set by analysis of variance (ANOVA) to ensure it had dose dependent response data suitable for mathematical dose-response modeling. Additionally, a post-assessment of models with acceptably fitting dose-response models should be applied to ensure models used going forward adequately account for the experimental data (Phillips et al., 2019; Thomas et al., 2007). This assessment utilizes minimum goodness of fit statistics to assess model fit, controlling for the 95% confidence interval (CI) of the BMD estimate and avoiding extrapolation of models beyond the range of the experimental data. Criteria for determining which genes amongst those with final acceptably fitted models should be used for ontology enrichment is the next step in ultimately deriving POD values. The intent is to have confidence that genes being used for enrichment are those with altered levels of expression as a direct result of cells being exposed to the chemical of interest. One such test for suitable genes proposed is the Williams Trend Test (WTT), which is a well-established metric for determining the presence of a monotonic dose dependent response (Williams, 1971, 1972). As an alternative, we propose using an established differential gene expression metric that is intended to capture not only the overall dose dependent response, but to characterize the specific gene changes at each concentration across the range of dose-response. In our analyses we have used DESeq2 for RNA-Seq and Biospyder TempO-Seq gene count data (Love et al., 2014). But there are numerous alternative approaches to differential gene expression analyses of gene count data that could also be applied (Liu et al., 2021; Robinson et al., 2009).

Once a gene set has been analyzed for ontology enrichment, appropriate methods to summarize enriched pathways to a single value for use as a POD must be applied. There have been numerous suggestions of how this should be accomplished but one emerging consistency when the POD are based on ontology pathway analysis is that often the summary methods differ by very little if the data used at each step in the analyses was selected with appropriate criteria of sufficient rigor (Farmahin et al., 2017; LaRocca et al., 2020b; Nault et al., 2020). In the analyses presented here we wished to assess the consistency of POD results when differing metrics for gene list selection are applied.

Data for a diverse group of industrial chemicals was recently published and thus available for analyses (Gwinn et al., 2020). We compare the tools already available in the BMDExpress2 program, ANOVA and the WTT, with DESeq2, a metric designed specifically for differential expression analyses of next-generation sequence count data (Love et al., 2014; Phillips et al., 2019). In contrast to these, we also selected genes by a magnitude of change metric using the absolute value of fold change greater than 2-fold. ANOVA and the WTT selected genes with a dose dependent response across all doses using a statistical significance of FDR<0.05 and absolute value of fold change greater than 1.5-fold (|FC|>1.5). DESeq2 was used to select the union of all genes significant by these same criteria but at any individual dose versus control contrasts. These four gene lists for each chemical were used to perform ontology enrichment analyses using the publicly available Reactome ontology (Jassal et al., 2020). The ontology enrichment was used to derive summary POD values for comparison across the four gene selection metrics to determine what issues may arise based on the metric for gene selection in a transcriptomic ontology pathway POD analysis.

## Methods

The chemicals selected (Table 1) were chosen from publicly available data and represent a broad range of MOA and industrial chemical classes. Each had been tested at 8 or 9 concentrations in 5-day in vivo rat exposures with 4 animals used at each concentration for biological replication (Gwinn et al., 2020). Samples from both liver and kidney had been analyzed using BioSpyder’s S1500+ Rat TempO-Seq platform. These data are available from the NCBI’s GEO database (accession GSE147072) and consist of a table of TempO-Seq RNA-seq counts for each of the 2653 probes included in the rat S1500+ transcriptomic screening panel. We only used samples with more than 2 million reads counted and with a Spearman’s rank correlation with its concentration cohorts of not less than 85%.

**Table 1.** Chemicals from 5-day rat exposures analyzed for differential gene expression and benchmark dose analyses.

| Chemical | Abbreviation | CAS number | MOA | Chemical use | Log<sub>10</sub><br>Max. Dose |
| --- | --- | --- | --- | --- | --- |
| Acrylamide | ACR | 79-06-1 | Neurotoxicity (interference with kinesin proteins) | Chemical production precursor (e.g. polyacrylamides) | 1.0 |
| Bromodichloroacetic acid | BDCA | 71133-14-7 | Tumorigenicity, cytotoxicity | Byproduct of drinking water treatments | 2.2 |
| Di(2-ethylhexyl) phthalate | DEHP | 117-81-7 | Endocrine disruption | Plasticizer (e.g. intravenous tubing) | 3.0 |
| Pentabromodiphenyl ether mixture | DE-71 | 32534-81-9 | Tumorigenicity (oxidative damage, possible epigenetic changes) | Flame retardant | 2.7 |
| Perfluorooctanoic acid | PFOA | 335-67-1 | PPARα agonist, oxidative stress, endocrine disruption | surfactant | 1.3 |
| 3,3',4,4'-Tetrachloroazobenzene | TCAB | 14047-09-7 | Ah receptor activity, tumorigenicity, endocrine disruption | Contaminant (pesticide production) | 2.6 |

### Differential gene expression analyses

Our intent was to examine the effects of criteria for selecting genes to be used in ontology enrichment analyses to derive significantly enriched pathways for computation of pathway-based POD values. The open source statistical program R was used to run the DESeq2 library (Love et al., 2014), and normalized log2(CPM+1) transformed data for BMD analyses was generated with DESeq2 for all analyses so that differing data normalization methods were not a factor in our comparisons. There are several methods for determining if a gene’s expression level is altered due to exposure to a compound (relative to unexposed controls) and these may yield different gene lists for input to an ontology enrichment analyses. Significant differentially expressed genes were determined using four metrics as shown in Figure 1: ANOVA, WTT, DESeq2, and a fold change filter for magnitude of change at treated concentrations relative to untreated vehicle control samples (Fisher, 1918; Fisher, 1919; Love et al., 2014; Williams, 1971, 1972). Both ANOVA and the WTT are methods of estimated the significance of a dose dependent response to exposure to a chemical and have been implemented in the BMDExpress2 analyses program (Phillips et al., 2019). DESeq2 is a library for the open-source statistical language R and uses a Wald test of dose specific contrasts for each concentrations response to exposure, relative to the untreated vehicle control samples. DESeq2 can thus test for an overall dose dependent response, as well as significance of response at each individual concentration used in the experimental exposure by simply creating the contrasts amongst the samples to be analyzed. In order to capture genes altered by exposure to compound with high confidence, we simultaneously applied both a statistical threshold of a false discover rate corrected p-value of less than 0.05 (FDR<0.05) and a magnitude of change threshold using an absolute value of fold change greater than 1.5 fold (|FC|>1.5) (Benjamini and Hochberg, 1995). However, with DESeq2 we selected the union of all genes which met these dual significance criteria for any dose from individual dose contrasts and not the overall dose dependent response. As an independent basis for comparison of post-hoc gene sampling, we also used a magnitude of change threshold only of a fold change of greater than 2-fold (|FC|>2). The value of 2-fold was used to reduce artifacts of noisy data when using magnitude of change in expression only, where gene lists at high fold change as a result of high dose exposure can be very large and include many non-specific genes (geness with no underlying ontology enrichment). The WTT and one-way ANOVA are built into the benchmark dose software BMDExpress2 (Phillips et al., 2019).

**Figure 1.**
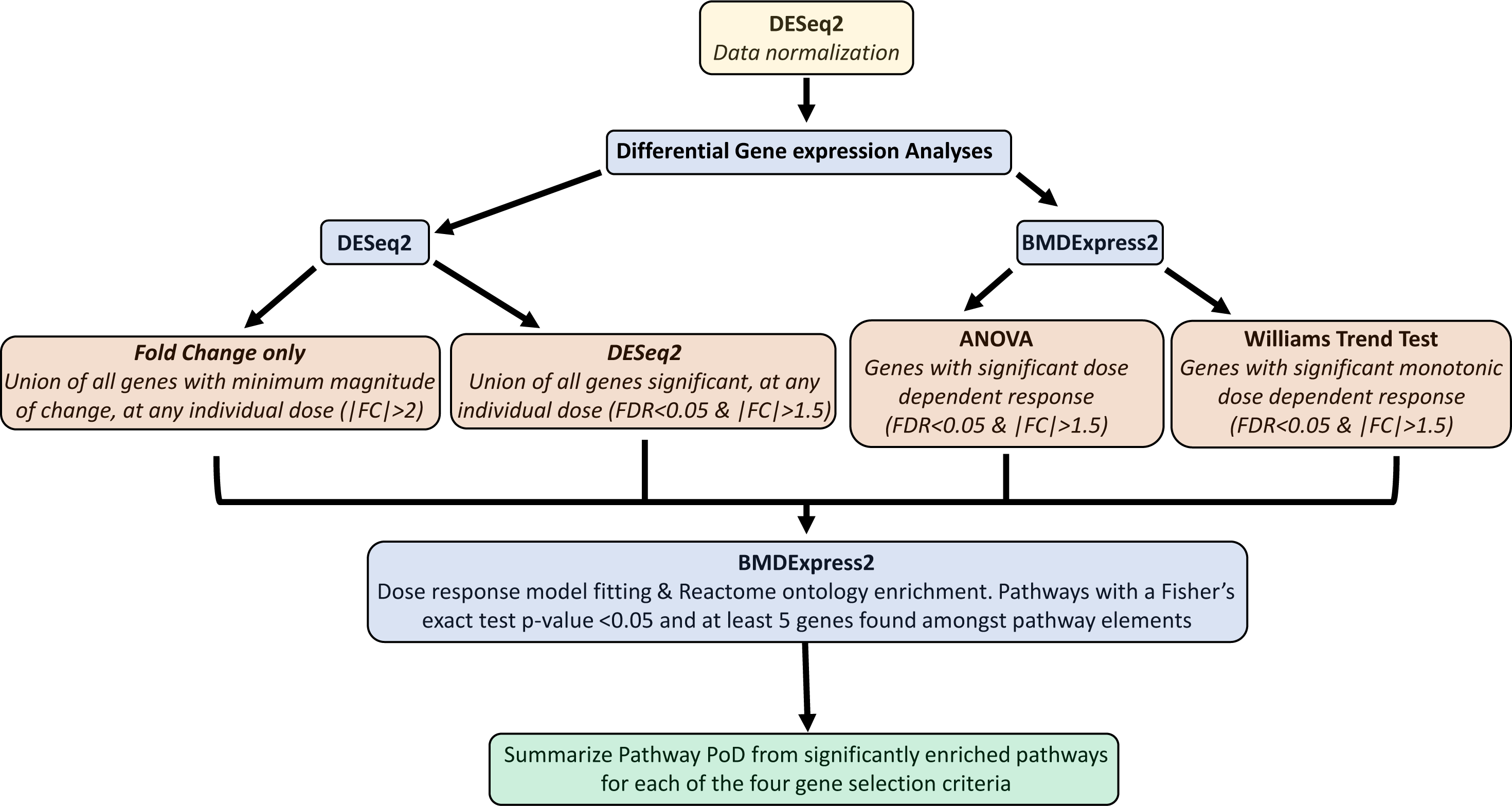
Workflow for analyses to derive ontology pathway-based POD values using 4 different criteria for selecting genes to be used for ontology enrichment analyses. All normalized data for BMDExpress2 model fitting was generated in DESeq2. DESeq2 was also used for differential gene expression analysis to determine significant changes in gene expression at each individual dose. BMDExpress2 was used to perform dose dependent statistical tests using one-way ANOVA and the William’s Trend Test. All dose-response model fitting and ontology enrichment performed in BMDExpress2.

### Benchmark dose modeling

We used the BMDExpress2 package for BMD model fitting and ontology enrichment (Phillips et al., 2019). A total of seven models were fit to genes: hill, linear, power, 2^nd^ and 3^rd^ order polynomials and exponentials. When fitting the hill model, the ‘k’ parameter was flagged if it was less than 1/3^rd^ of the minimum concentration used in the exposures, and the next best fitting model was selected instead based on p-value (<0.05). Power models were restricted to power coefficients of 1.0 or greater. A benchmark response (BMR) value of 1 standard deviation was used for BMD calculations. All models were fit assuming constant variance as the data was log2 transformed after the normalization in DESeq2. A final best fitting model was determined by first determining the best fitting polynomial model by nested Chi square test. The best fit polynomial model was then compared to the remaining models to select a final overall best fitting model by Akaike information criteria (AIC) (Akaike, 1998). Once best fitting models were selected for each probe in the rat S1500+ panel, three quality control (QC) thresholds were applied to select the final genes with acceptable BMD model fits. First, any model with extrapolated BMD value beyond the highest concentration used in the dose-response experiment was rejected. Similarly, any best fitting model with a goodness of fit p-value less than 0.1 was rejected. With the likelihood ratio test performed on the models, a larger p-value indicates a better fit between the model and the experimental data (Thomas et al., 2007). Finally, any model which had a BMDU/BMDL ratio greater than 40 was rejected to avoid BMD estimates with excessively large 95% confidence intervals.

The remaining genes were then sorted by the gene lists derived from the four metrics for significant differential expression to prepare final gene lists with good fitting dose-response models for ontology enrichment. Ontology enrichment was performed in BMDExpress2 program using the publicly available Reactome ontology (Fabregat et al., 2018; Jassal et al., 2020). BMDExpress2 performs a conventional over-representation test of the query gene lists relative to the pathway annotated gene lists in Reactome. We set a significance threshold for enrichment of a pathway having at least 5 of our query genes found amongst the pathway elements, and with a Fisher’s exact test p-value from the over-representation test that was less than 0.05. A pathways BMD value is computed as the median gene based BMDU/BMD/BMDL values for the genes found amongst the pathway category elements.

For ontology enrichment results where more than 20 Reactome pathways were significantly enriched (more than 5 elements found amongst the pathway elements, and a Fisher’s exact test p-value < 0.05) we summarized average pathway median BMD values using five approaches:

- 20 most statistically significant as ranked by Fisher’s exact test p-value
- 20 pathways with lowest median BMD1SD
- 20 pathways with the most elements (i.e., most query genes found amongst pathway elements)
- 20 pathways with most altered genes as ranked by maximum absolute value of fold change (|FC|)
- Average (median values) for all enriched pathways

When less than 20 pathways were significantly enriched, we modified the summary methods to use instead:

- The pathway with lowest Fisher’s exact test p-value
- The pathway with lowest median BMD1SD
- The pathway with the most elements
- The pathway with most altered genes by |FC|
- Average (median values) for all enriched pathways

Comparisons were restricted to median pathway values or average median pathway values as our experience with other data (not shown) has shown these are consistently more conservative that pathway mean values.

## Results

Table 2 lists the differential gene results for the four metrics used. The numbers shown for each chemical and metric are the number of genes which met the significance criteria of FDR<0.05 and |FC|>1.5 fold. While the DESeq2 analyses determined significantly differentially expressed genes with all chemicals, ANOVA and WTT did not in several instances. These analyses test for significance change across the entirety of the dose range to determine overall dose-dependent changes. While DESeq2 can also analyze the data in this same way, it also allows examination of discrete changes at each dose as was done here, and thus may detect changes in gene expression that are not strictly part of a dose dependent trend across all doses. The large number of altered genes by fold change alone, but which do not pass a statistical threshold of significance is a common observation in gene expression data due to the inherent variability of gene expression (Rapaport et al., 2015).

**Table 2.**
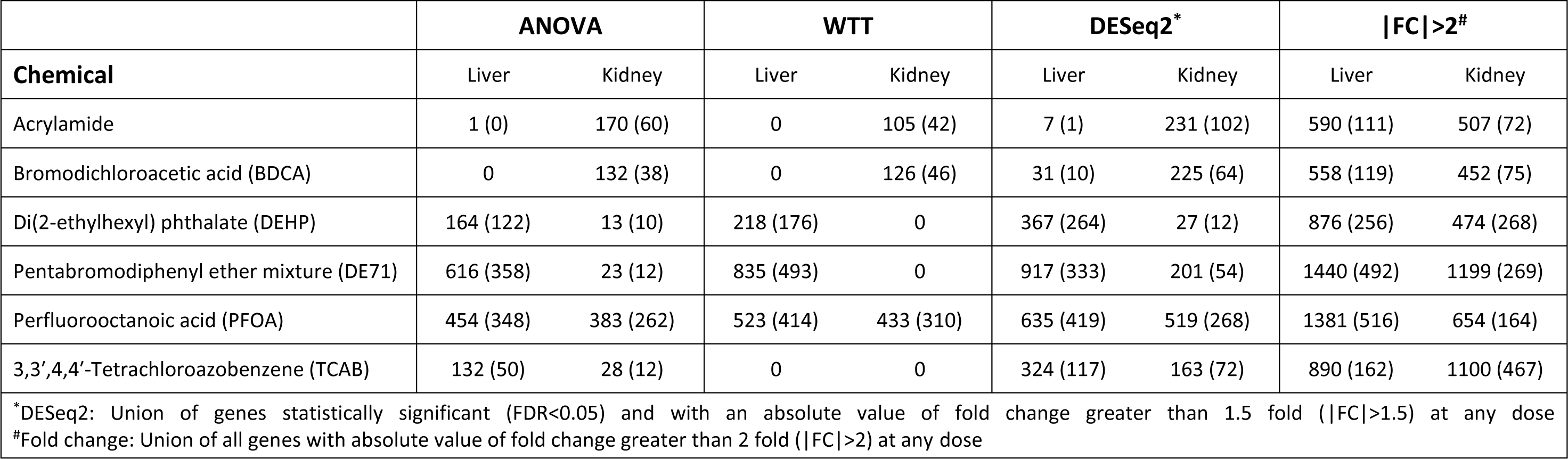
The number of genes significant by a false discovery rate corrected p-value (FDR<0.05) and with a magnitude of change of greater than 1.5- fold using different proposed metrics for significance. In parentheses are the number of those significant genes with a passing best fit model after BMD analyses. A best fit model must have a BMD less than the maximum treatment dose, a goodness of fit p-value greater than 0.1 and a BMDU/BMDL ratio less than 40.

In parentheses are numbers of significantly differentially expressed genes that had a best fit BMD model that met the post model-fitting QC thresholds (overlap in these genes for kidney in supplemental figure 1, and for liver in supplemental figure 2). These represent genes suitable for ontology enrichment and subsequent ontology pathway derived POD derivation. One chemical in liver, BDCA, failed to produce any final gene results with ANOVA due to a lack of statistical significance, despite many genes with a large magnitude of change. In the case of ACR in liver, the one gene that met our criteria for significant differential gene expression did not have an acceptable best fit BMD model (the goodness of fit p-value was <<0.1). Similar to the outcome of ANOVA tests with the compound BDCA, with the Williams Trend Test the issue with two compounds—ACR and BDCA—was a lack of statistical significance, not an issue with magnitude of change in genes. This was also true with the WTT and DEHP and DE-71 in kidney samples, and TCAB in both liver and kidney samples. In contrast to ANOVA and the WTT, our analyses in DEseq2 is designed to test for changes in gene expression at discrete doses, regardless of the nature of the overall dose-response. Subsequently, every one of these chemicals have some significantly differentially expressed genes, even though many of these do not fit a dose-response model well (as shown by the numbers in parentheses). The application of a simple fold change threshold produces the largest number of differentially expressed genes, but historically such a limited significance threshold often produces many genes that are likely to be a consequence of cellular changes and not part of the cellular functional response mounted by cells to deal with the perturbation of chemical exposure (Jong et al., 2019). As shown in Table 2, many of those most altered genes fail to produce best fitting BMD models that meet QC assurance of a well modeled dose-response.

Gene expression changes by themselves don’t provide inherent details of cellular processes and mode of action. Numerous methods of associating gene based changes with known functional cellular biology have been developed (Alexander-Dann et al., 2018; Barel and Herwig, 2018). BMDExpress2 uses a Fisher’s exact test in an over-representation ontology enrichment analysis to determine if any overlap in the gene list provided with the pathway elements is unlikely to have occurred by chance. Any set of genes found amongst the pathway elements can be considered a biologically relevant gene cohort, with their gene expression profile being altered due to exposure to the test compound. This method of enrichment filters gene lists by know functional elements in cellular biology. By determining which cellular functions are likely affected by changes in gene expression, an hypotheses of the cellular mode of action in response to exposure to a chemical can be derived. By using this same approach when deriving a POD value, the pathways used in summarizing that POD value should be functionally relevant to the elements of the MOA.

After ontology enrichment we found numerous enriched Reactome pathways for some compounds, and none or few for others (Table 3) reflective of the numbers of DEGs available for enrichment analyses. The distribution of enriched pathways for PFOA (supplemental figure 3) from kidney samples exposed to PFOA were then summarized to single values (PoDs) and plotted in Figure 2. In transcriptomic BMD analyses, an enriched pathway has a BMD value derived from the best fitting model BMD value for each gene found amongst the pathway elements. A POD can then be derived for any set of pathways by summarizing the individual pathway values, using any one of several proposed summary methods (Farmahin et al., 2017; Izadi et al., 2012). With PFOA, genes significant by ANOVA produced the most conservative estimates for BMDU, BMD1SD and BMDL, but was also very similar to the values produced from genes significant by the WTT. DESeq2 and genes with |FC|>2 produces larger estimates for all values with those for genes with |FC|>2 being much higher and with large 95% confidence intervals (CI). BMDL values for DESeq2 significant genes were on average close to twice that of those for either ANOVA or the WTT.

**Figure 2.**
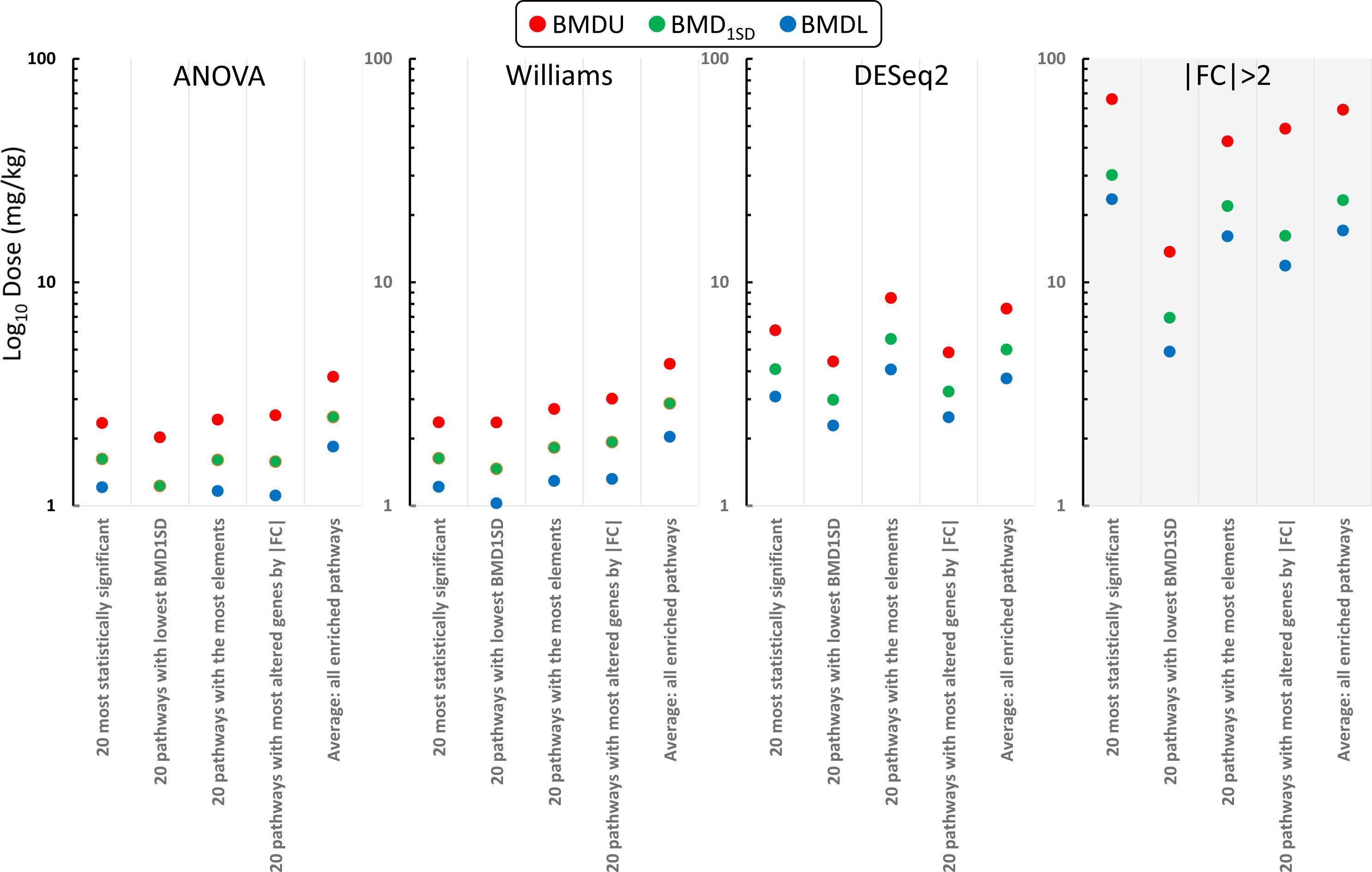
Pathway BMD summaries for PFOA in kidney. Pathways enriched with genes selected by fold change only nor only have the highest summary values, but also the largest 95% confidence intervals. Pathways enriched with genes detected by ANOVA and the Williams Trend Test were highly congruent and more conservative than those from DESeq2 gene enriched pathways.

**Table 3.**
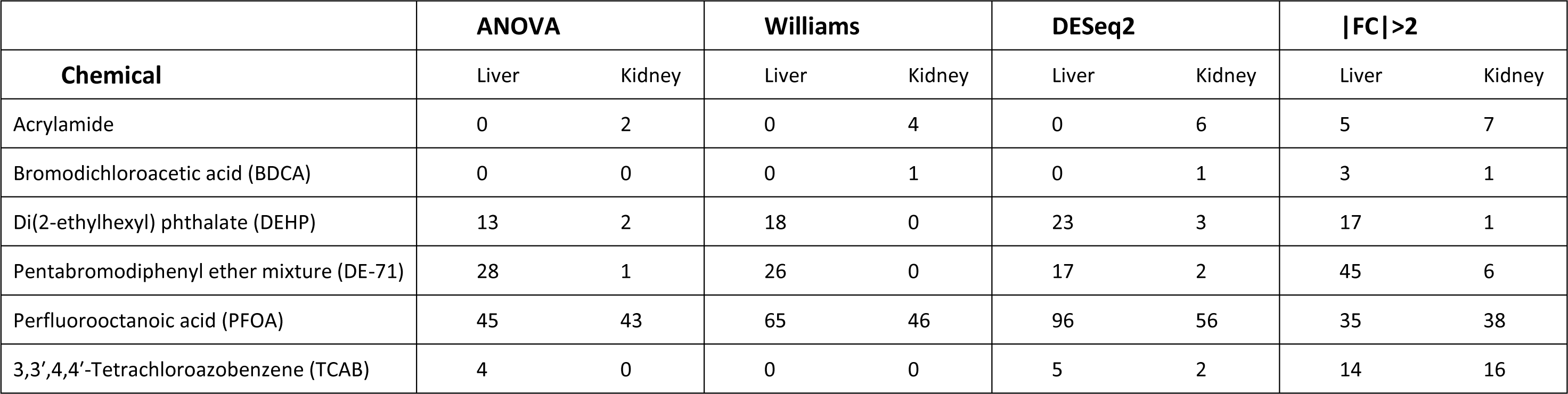
The number of significantly enriched Reactome pathways from enrichment analyses of the genes both significant by one of four metrics and passing best fitting model QC metrics.

Figure 3 plots summary values for PFOA from liver samples, and values using the three statistical test metrics are all very similar to each other, although in liver, the WTT gene set produced the most conservative BMD and BMDL summary values. Values for DESeq2 were higher than for ANOVA or the WTT but by a much smaller margin than seen in the kidney samples. Once again, genes significant only by a fold change threshold produced summary values typically larger than those generated by the three other metrics.

**Figure 3.**
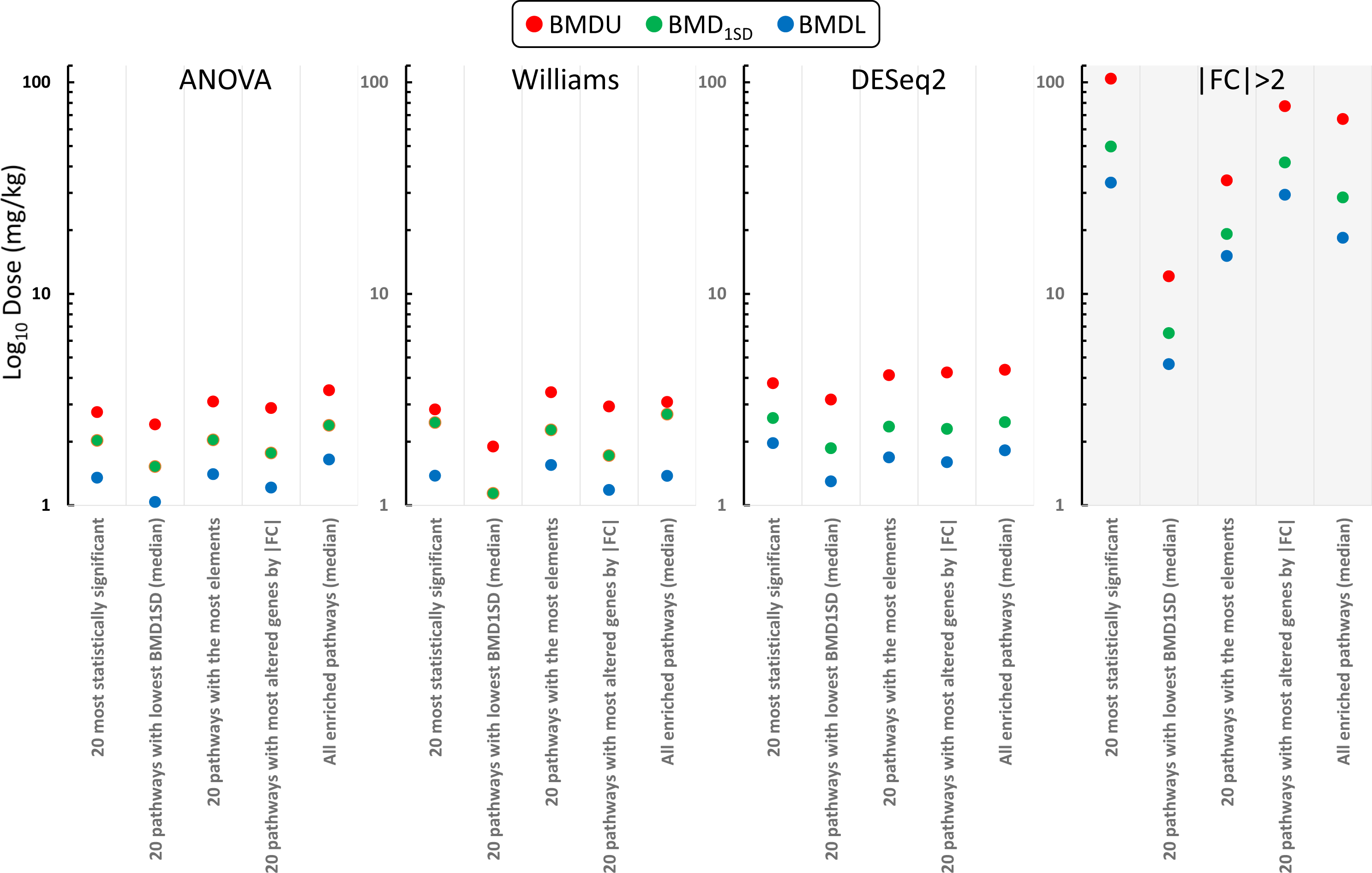
Pathway BMD summaries for PFOA in liver. As seen in the kidney samples, pathways enriched with genes significant by fold change only are the least conservative. Pathway POD values for the three statistical metrics are consistent. The Williams Trend Test had the most conservative BMDL value (20 pathways with lowest BMD1SD value) of 0.73 mg/kg.

Pathway distribution for significant pathways for DE-71 exposure in liver samples (supplemental figure 4). also produced some enriched pathways uniquely significant with each metric, but with a core set of common pathways. The ANOVA significant pathways were the most sensitive using the average median value of the 20 pathways with lowest BMD1SD value, although this same summary from the |FC|>2 group was very similar (Figure 4). DESeq2 significant genes resulted in only 17 significantly enriched pathways so these are summarized by their overall average median values as well as select single pathways using the most statistically significant pathway, that with the lowest BMD1SD median value, the pathway with the most elements, and the pathway with the most altered genes as measured by maximum absolute value of fold change. By these criteria, three of them (pathway with lowest BMD1SD, pathway with most elements and pathway with most altered genes) are not greatly different from the |FC|>2 summary values or the WTT values and only the ANOVA values were consistently more conservative by a small margin.

**Figure 4.**
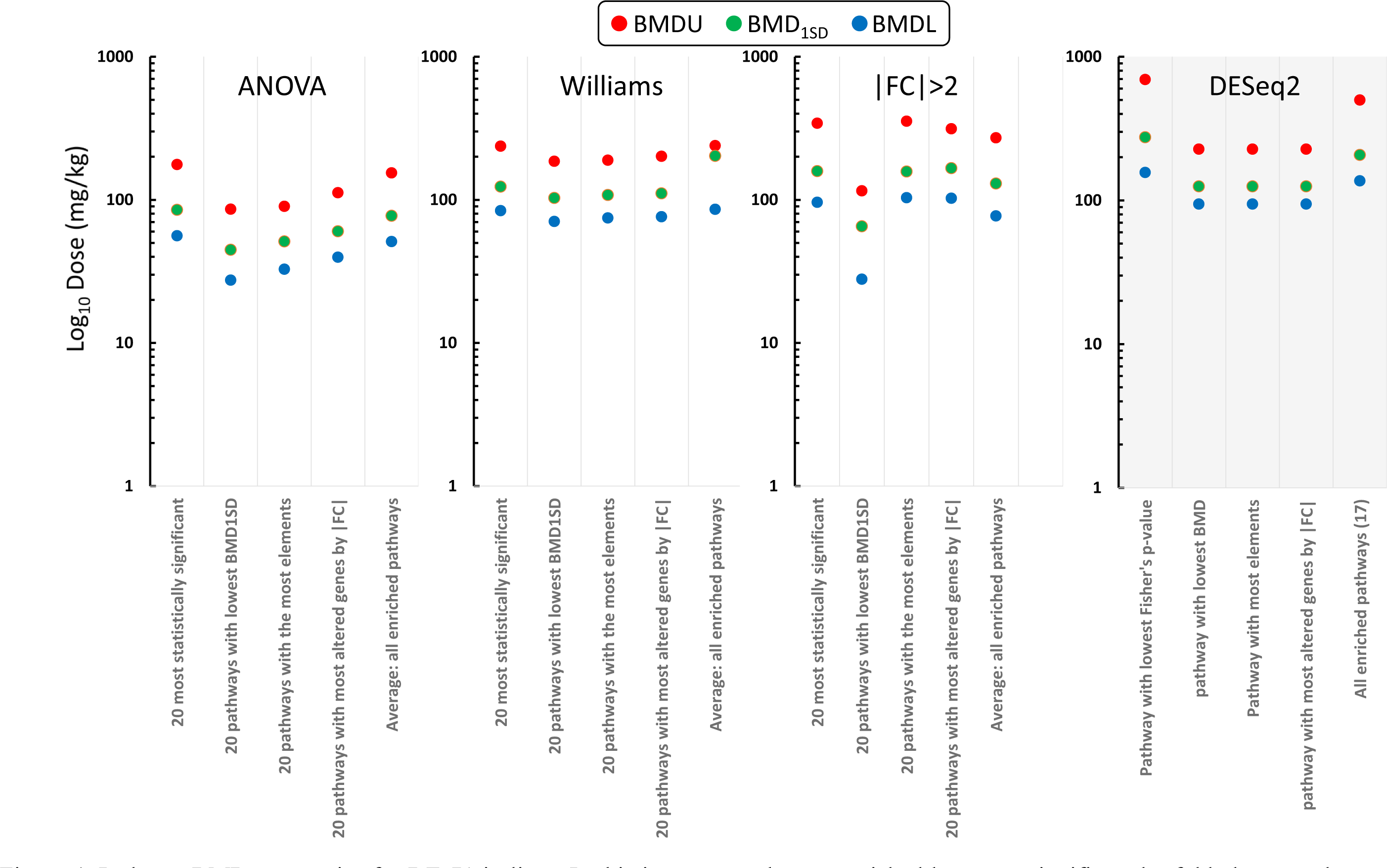
Pathway BMD summaries for DE-71 in liver. In this instance, pathways enriched by genes significant by fold change only were not greatly different from the other three metrics, and produced the second most conservative BMDL (27.94 mg/kg). Pathways enriched with genes significant by ANOVA had the lowest BMDL overall (27.47 mg/kg), but the differences in values across all four metrics was not greatly different.

The final comprehensive comparison of summary values possible was that for DEHP in liver samples (supplemental figure 5). Differences amongst the gene sets used reflect differences in downstream child processes significantly enriched with the different gene sets, focused on a common core set of parent categories enriched with all. Only one gene set, DESeq2, resulted in more than 20 enriched pathways and so only it allowed for the preferred summary method of averaging 20 pathways ranked by different criteria. It and the other three metric’s summary values were instead summarized as with DESeq2 was with DE-71. These data are plotted in Figure 5. The individual pathway values show very little deviation for BMDL and BMD1SD which reflects the overlap in pathways detected with 11 pathways significantly enriched using genes from all four metrics. With DESeq2 there were 23 enriched pathways, and the summary using 20 pathways is shown at the far right of Figure 5. In this case, DESeq2 also has 7 significantly enriched pathways not detected at all with the other three metrics, and its pathway average summary values reflect the input of these additional pathway values resulting in less conservative values for BMDL and BMD1SD, as well as larger 95% CI.

**Figure 5.**
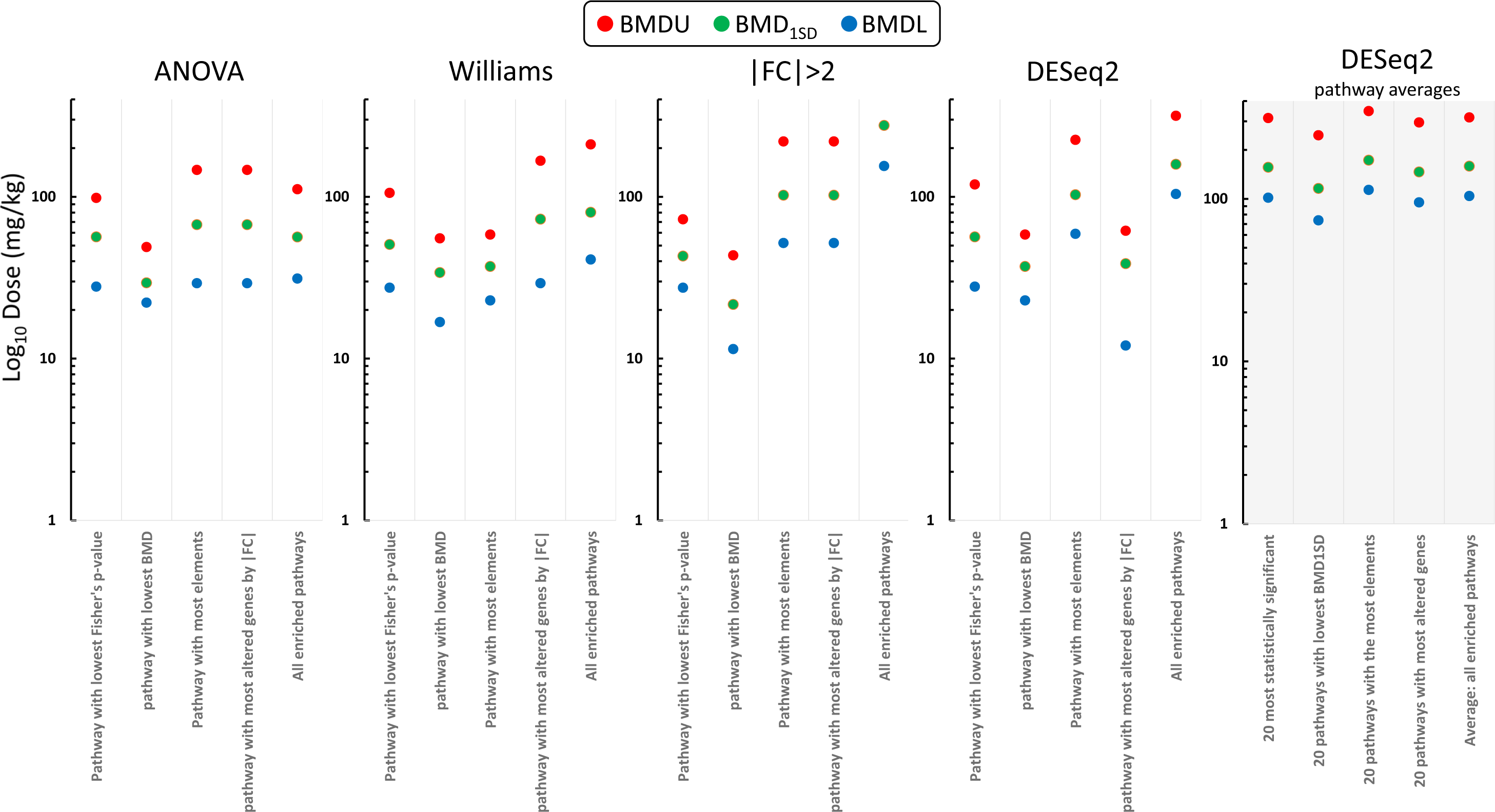
Pathway BMD summaries for DEHP in liver. As only genes enriched by DESeq2 produced more than 20 enriched pathways, it was the only result summarized by our normal summary metrics and this is shown on the far right plot. Beside this is the summary by individual pathways in order to provide direct comparison to the other three metrics where fewer than 20 pathways were enriched. The individual pathways enriched with genes significant by ANOVA and the Williams Trend Test do not differ greatly and are slightly more conserved that the multiple pathway summaries by DESeq2. The most conservative BMDL value was derived with genes significant by fold change only, where the pathway with lowest BMD1SD had a BMDL value of 11.45 mg/kg. This compares to that derived from the single pathway with the most altered genes by |FC| with DESeq2 significant genes, where the BMDL was 12.03 mg/kg.

Table 4 summarizes the pathway enrichment for those instances where only a single or few pathways were significantly enriched. Acrylamide kidney samples produced similar values using any of the four metrics, with a simple fold change filter being the least conservative, and DESeq2 the most conservative based on the average of 6 enriched pathways. Similarly, the WTT and DESeq2 produces similar values and for the same single significantly enriched pathway for BDCA, however DESeq2 had 7 elements in this pathway contributing to its pathway value while the WTT only had 6 genes in the enriched pathway (the two analyses had 5 significantly differentially expressed genes in common for this pathway). DEHP in kidney shows the least consistent summary values for three of the four metrics with significantly differentially expressed genes and again the differences in values are reflective of the genes that were part of the set used for over-representation analysis. Values for DE- 71 in kidney were similar for ANOVA and DESeq2 but these two metrics also resulted in few genes for enrichment (12 with ANOVA and 54 for DESeq2). TCAB in Liver is the final comparison in Table 4 and it also shows considerable variation in summary values for BMDL and BMD1SD, reflecting differences in pathway elements for common pathways as well as some differences in enriched pathways found.

**Table 4.**
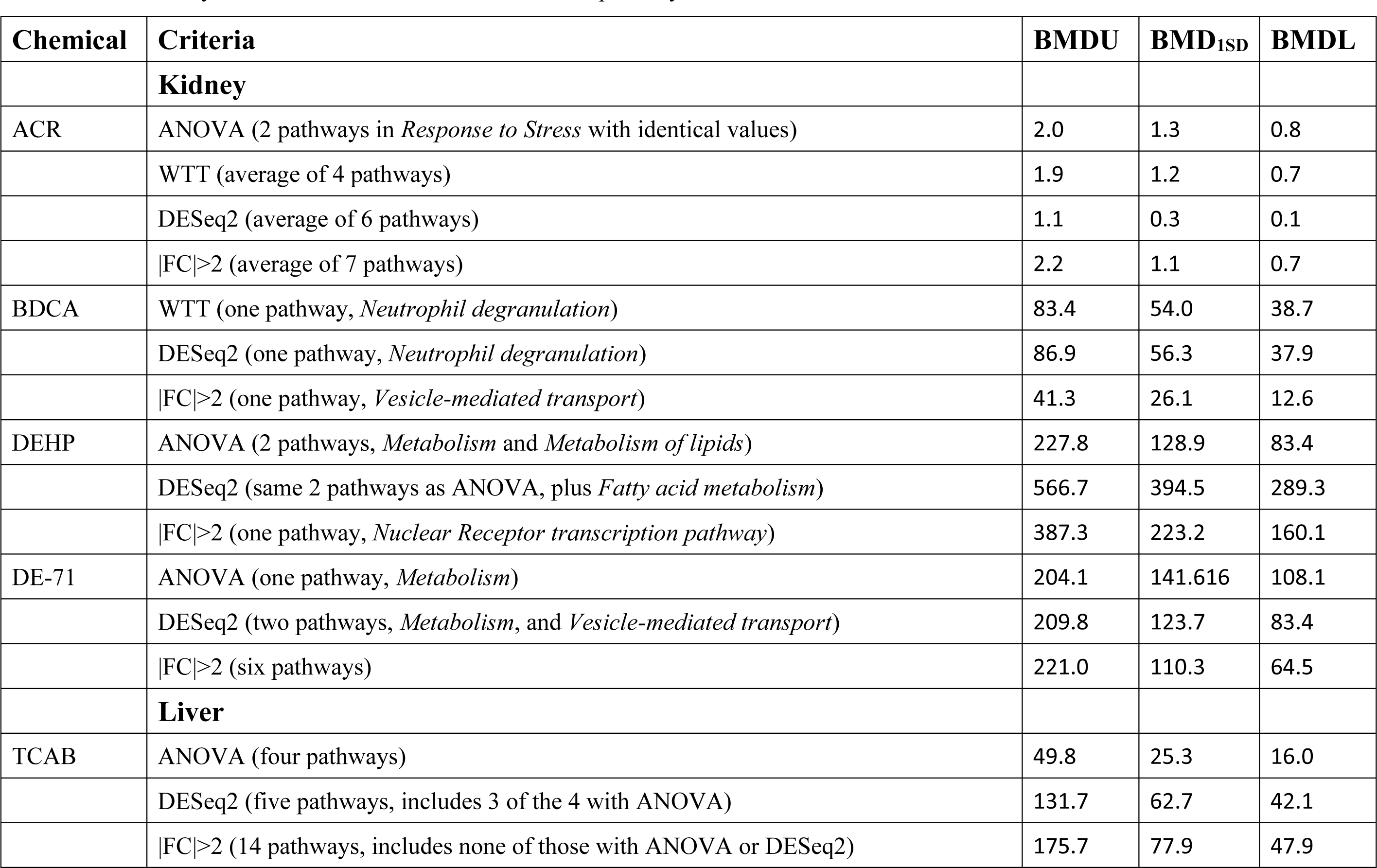
Pathway values for instances with few enriched pathways.

## Discussion

Transcriptomics has become a widely used tool for inference of mechanisms of cellular biological response upon exposure to a chemical. By examining changes in gene expression when cells are exposed to a chemical, relative to comparable cells from unexposed samples, key events in the progression of functional changes in cells can be identified and tracked over a range of doses as well as over time for chronic exposures. This has become a highly useful method to understand the development and progression of adverse response to chemical exposure. As changes in gene expression precede observable apical effects, differential gene expression analyses also provide a mechanism to infer functional response to chemical exposure from short term exposures, as well as the use of in vitro exposures in place of animal studies.

Benchmark dose analyses was adapted for transcriptomics data over a decade ago and combined with cellular biology ontology analyses to use BMD dose-response models in the analyses of functional relevant response to exposure to chemicals (Thomas et al., 2007; Yang et al., 2007). This analytical technique was then used to demonstrate short term transcriptomic changes correlated with apical endpoints and thus provide an opportunity to develop methods for rapid assessment of chemical hazard from short term dose-response studies (Thomas et al., 2013b). Interest in the use of transcriptomic BMD analyses to derive POD values relevant to chemical hazard screening and decision making has grown steadily as both a cost and time saving approach as well as an important component of moving away from in vivo to in vitro chemical screening. In order to derive meaningful POD values for hazard assessment, the value of transcriptomics is the ability to combine BMD analyses with functional ontology analyses to anchor potential POD values in the functional response of cells to exposure of the chemical in question. However, dose-response to different compounds in different tissues or cells can be highly variable. This raises the question of how to design best practices to ensure that the analyses leading to a proposed POD value are the most suitable for a given set of exposure data.

Early in the development of using transcriptomic data in toxicology, the FDA led a coordinated effort to develop proposed best practices for consistent generation and analyses of toxicogenomic data from microarray technologies (the MicroArray Quality Control consortium). One of the tenants derived from this effort was the application of simultaneously using both a statistical criterion and a minimum magnitude of change when determining significant changes in gene expression in treated samples relative to untreated control (Guo et al., 2006). The rational for such a best practice was based on the observations that alternate analyses of data from different sources or generated at different times varied in terms of the specific genes displaying significantly altered expression profiles. However, when this dual significance criterion was applied, the aggregate gene sets with significant changes in their expression produced very consistent functional ontology enrichment results, allowing for consistent cellular MOA inferences from alternate data sets (i.e. different dose-response exposures in time or place or different platforms in the detection of levels of expression) for a given chemical.

Dose response experiments analyzed using this approach have been useful in understanding the cellular effects of exposure to a diverse set of chemicals (Alexander-Dann et al., 2018; Andersen et al., 2017; Andersen et al., 2018a; Black et al., 2012; Black et al., 2015; Efremenko et al., 2014; McMullen et al., 2019; Nault et al., 2020; Thomas et al., 2013b). Of interest then is do these same principles apply when combined with BMD analyses and ontology enrichment to use significantly enriched ontology pathways to derive summary BMD and BMDL values for use as POD values in chemical screening and assessing hazard? There are multiple statistical approaches for determining significance of change in gene expression due to exposure to a chemical, and how does the choice of method affect the outputs from a BMD ontology enrichment analysis? Common methods for analyzing gene expression data for differential expression include ANOVA, Bayesian methods, General Linear Models and Wald tests, and numerous non-parametric approaches (Breitling et al., 2004; Churchill, 2004; Kendziorski et al., 2003; Ritchie et al., 2015; Townsend and Hartl, 2002).

Tools such as ANOVA and the WTT are useful for detecting a dose dependent response in a multiple exposure experiment, but they do not address the question of what specific differences exist between any given treatment dose and the untreated control samples. It is the latter that is most frequently applied in transcriptomic MOA studies in order to detect the progression of cellular events as dose increases. ANOVA can be combined with post-hoc contrasts to determine discrete significant differences for each dose group but that was not done in the present analyses. This consideration is important in terms of determining which genes are most indicative of the underlying cellular functional response, and which genes are most suitable for use in ontology analyses for summary to a single derived POD value. Ideally, a transcriptomic POD value would be based on genes both anchored in the cellular biology of the response to a chemical and also which have a well-defined dose-response reflective of the sensitivity of the response. By this we mean identifying genes anchored in the cellular biology of the response, displaying changes in expression specifically due to chemical exposure that can be identified with high confidence. And a well-defined dose-response for a gene means a dose-response where the final fitted mathematical dose-response model summarizes the experimental data with high confidence. Transcriptional dose-response data is inherently noisy data as expression levels constantly fluctuate in any random population sample of cells, and it is not clear yet how specific metrics for selecting genes for POD derivation may affect the outcome given how varied transcriptomic dose-response results can be.

We examined just four possible metrics for significance of differential expression amongst several that could be used, and as expected the specific gene sets identified as displaying significant differences in expression due to chemical exposure varied amongst these four. Applying a simple fold change only generated many genes which were not statistically significant by the three metrics we used (ANOVA, WTT and DESeq2), with the exception of PFOA exposure in liver samples. Given that all 6 of the compounds examined are known toxicants, it is not unexpected that there would be large gene expression changes that would not be detected as statistical significance as adverse response in cells usually produces some gene changes that are simply a consequence of cellular stress or a result of functional response and not part of the coordinated functional response itself. Changes in cell cycle for example may trigger changes in expression of DNA damage and replication pathways as normal DNA replication sentinel pathways are affected due to cell proliferation or suppression of cell cycle.

In the case of PFOA in liver, we examined the gene lists in the Reactome online analyses tool, using Reactome’s pipeline to map rat gene IDs to their human homologues to take advantage of the more extensively annotated human ontology. There was a core set (supplemental figure 4) of high order parental pathways detected with genes from all four gene sets (*Cell Cycle, Metabolism, Signal transduction, Immune System, Gene expression, Programmed cell death, Cellular response to external stimuli, Extracellular matrix organization, and Developmental biology*). The differences between the pathways were in the numbers of child processes detected downstream of these large parental pathway categories. In particular, all showed enrichment of some pathways in *Programmed cell death* but only genes determined as significant by fold change or significant by DESeq2 enriched two terminal pathways in *Programmed cell death* not seen in the enriched pathways with the other 2 gene lists (terminal child processes of *Caspase activation* via *Extrinsic apoptotic signaling pathway*). The gene for the Fas cell surface death receptor is significantly up-regulated at the highest dose of 20 mg/kg. These apoptotic cell pathways are indicative of some degree of cytotoxic response with PFOA at the highest doses of 10 and 20 mg/kg even if it had not progressed sufficiently after 5 days to manifest itself in any apical observed response. The original publication of these data did reference that terminal body weights were significantly reduced in rats exposed to PFOA, and liver weights were increased (Gwinn et al., 2020) This was also evident in the kidney PFOA samples, although the response does not seem as drastic as that displayed in liver and the terminal caspase activation pathways seen in liver were not enriched in kidney. This then in turn explains the enrichment summary for the |FC|>2 metric in Figures 4 and 5, where the PoDs derived with this gene set produced summary values larger than with the other three metrics, and with the largest 95% CI as a consequence of large-scale gene expression changes that were not statistically significant and are likely largely the response of a cytotoxic response at high dose. With the application of a statistical filter to remove those noisy genes with large fold change, the three statistical metrics resulted in very similar pathway POD values and consistent 95% CIs. Given the original experiment used 8 treatment concentrations, and evidence for cytotoxicity is only evident at the highest concentration, this would be a case where dropping the maximum treatment concentration from the BMD analyses may be preferred prior to attempting to derive pathway-based POD values.

The chemical DE-71 did not show any such drastic difference with the POD summary results for the fold change metric although this method produced insufficient enriched pathways for a robust pathway summary. Since most of the differentially expressed genes in this case were less than 2-fold different from untreated control samples, this would also indicate a less robust dose-response than seen with PFOA in liver. Similarly, with DEHP in liver, the three statistical metrics produced very similar ranges in possible POD values, while the fold change metric resulted in the least conservative estimates. However, only the DESeq2 analyses indicated sufficient differentially expressed genes that resulted in more than 20 enriched pathways, again indicating a weak dose-response in terms of functional information derived from the gene expression changes.

In general, the application of an appropriate statistical significance metric combined with a minimum magnitude of change threshold produces very consistent POD values from BMD pathway enrichment analyses. As previously observed, the specific method of summarizing pathway values to a single POD seems to have little effect on the derived values (Farmahin et al., 2017). A caveat to this would be designing a sufficiently robust dose-response experiment to ensure that sufficient response is induced to capture a biologically meaningful differential expression response. Confounding issues can be a maximum dose being too high, as apparent with PFOA in the data presented here, where cellular response to extreme stress and cytotoxicity can obfuscate the chemical specific cellular response to exposure and inflate or at least bias any derived POD values due to gene expression changes induced by extreme cellular stress or cells entering terminal apoptosis. Of the metrics used in these analyses, DESeq2, used a series of concentration specific contrasts and merging significant genes from any doses into a single list for each chemical, was able to detect significant gene changes for all 6 compounds. On the other hand, both ANOVA, and the WTT, failed to detect any significant changes in several cases. This reflects that while these test for an overall dose dependent response, they are not designed to detect significant changes for any specific concentration tested. The WTT in particular tends to indicate no significant dose-response when the nature of the dose-response is more of a threshold response than a monotonic dose-response, and this is a fundamental difference between testing simply for a trend across all data versus significant differences at specific tested levels (Hamada, 2018). An experimental dose-response may manifest as a threshold response if the compound truly exhibits threshold response behavior or because the dose range tested was not appropriate for capturing a non-threshold response. This explains why the WTT indicated no significantly altered genes with TCAB in either tissue. With DESeq2 we found that of the 324 genes with FDR<0.05 and |FC|>1.5, over 80% of those genes were only significant at one or both highest doses (200 and 400 mg/kg). Similarly, with DEHP in kidney samples, DESeq2 detected no significant genes at the lowest five doses, and 21 of the 27 significant at any dose were only significant at the two highest doses. For compounds with similar moderate response where the dose range used is not optimal to detect the influence of chemical on expression levels, the WTT usually will fail to detect any significant changes. ANOVA also failed to detect any significantly altered genes with BDCA in liver samples where again the response did not involve large numbers of genes. With DESeq2, BDCA liver samples had 31 genes significant at any dose, but 24 of those were only significant at the highest dose. An ANOVA analyses can always be modified to include post-hoc linear contrasts to provide an analysis more analogous to that of DESeq2 than a simple yes/no trend result.

Transcriptomic studies in toxicology are highly varied. Studies vary from the very large scale to individual chemical comparisons (Harrill et al., 2021; Nault et al., 2020; Rowlands et al., 2013). The focus may be on relative hazard screening, or detailed cellular MOA. Transcriptomic BMD-based POD analyses may be targeted to potential regulatory application, or for relative potency comparisons in a small group of lead compounds in development. Multiple methods of differential gene expression analyses are in common use and are chosen for a study based on the data requirements to meet the goals of a study. If, as we have shown, the choice of a particular metric for deriving significantly altered genes is not material, this allows tailoring of differential gene expression with BMD analyses and POD derivation to meet multiple research goals. Instead, methods can be built around the most appropriate for a given chemical dose-response to facilitate integrated analysis pipelines suitable for a wide range of applications. Future pipelines for BMD analyses could include tools to generate data to inform how those analyses should be concluded. As mentioned, the simple addition of post-hoc linear contrasts to an ANOVA would provide information on not only the nature of an overall dose dependent trend in the treated samples, but also which concentrations tested showed significant differences and provide output for detailed MOA analyses. Tools could be included for examination of maximum dose significant gene lists for detection of evidence of cytotoxicity to allow for decisions about removing high(est) doses from consideration of the functional response and POD derivation. A differential gene expression analyses has already recently been combined with BMD analyses in a single integrated analyses pipeline by researchers at McGill University using LIMMA for differential gene expression (Ewald et al., 2020). Independent BMD packages include the *bmd* library for the statistical language R also are available and can accommodate any data input from prior analyses (Jensen et al., 2020). Finally, the addition of tests for threshold response, and models specific for such dose-response would allow for more appropriate analyses of data where the dose-response is observed only on the high(est) doses used or where the nature of response to the chemical is truly a threshold response (Spassova, 2019).

## Acknowledgements

This work was supported by the American Chemistry Council’s Long-range Research Initiative.

---

**Supplemental figure 1.**
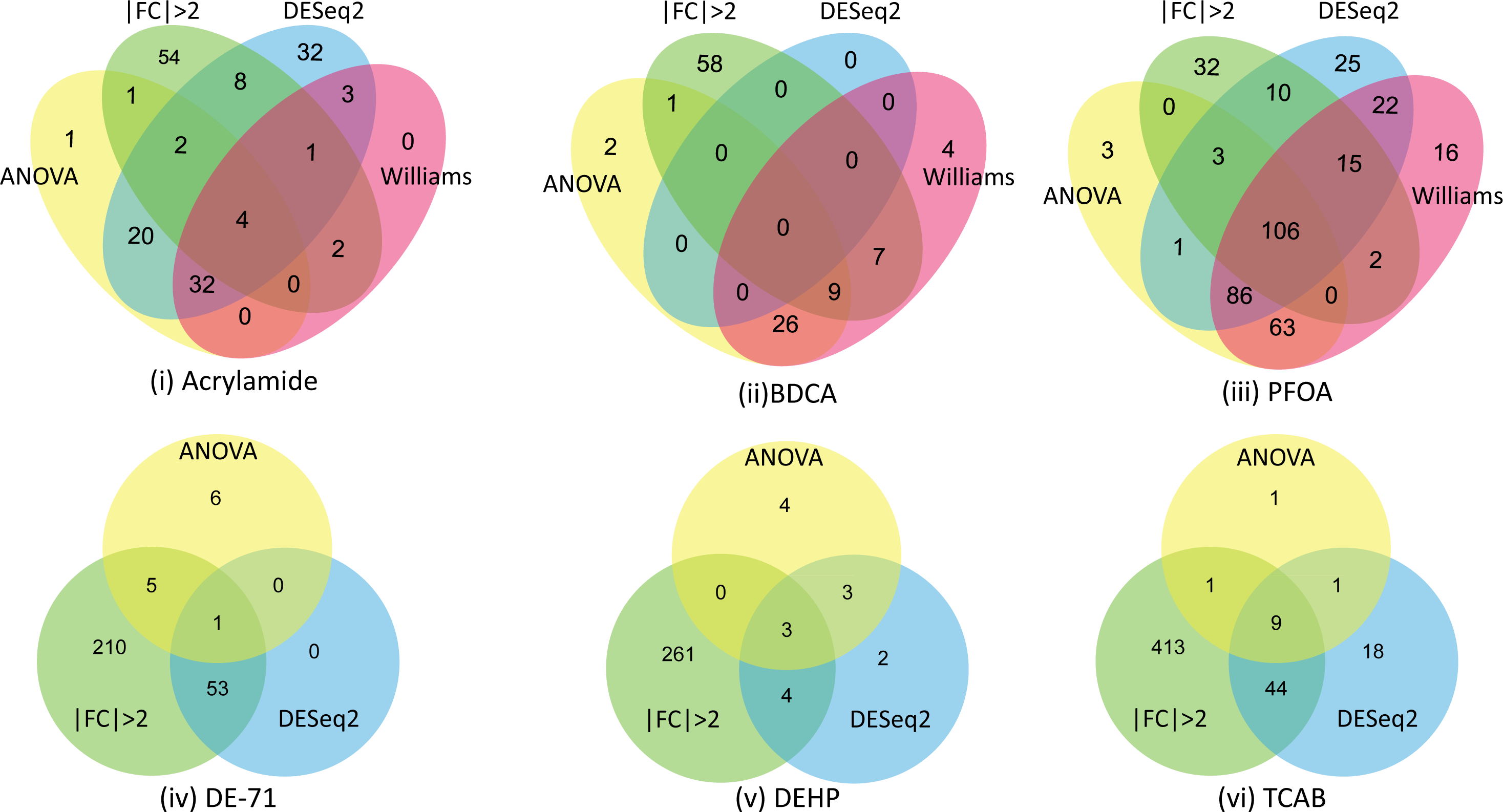
Distribution of significant genes (FDR<0.05 & |FC|>1.5) which also had a BMD best fitting model that passed model QC assessment (BMD <maximum experimental dose & goodness of fit p-value <0.1 & BMDU/BMDL <40) from in vivo rat kidney samples. The Venn diagrams in the lower row do not include any genes for the Williams Trend Test as no genes met these criteria. As indicated, genes significant by fold change alone have the most unique genes, as many such genes do not pass any statistical threshold. Even when a statistical metric was used, each metric often has numerous unique significant genes.

**Supplemental figure 2.**
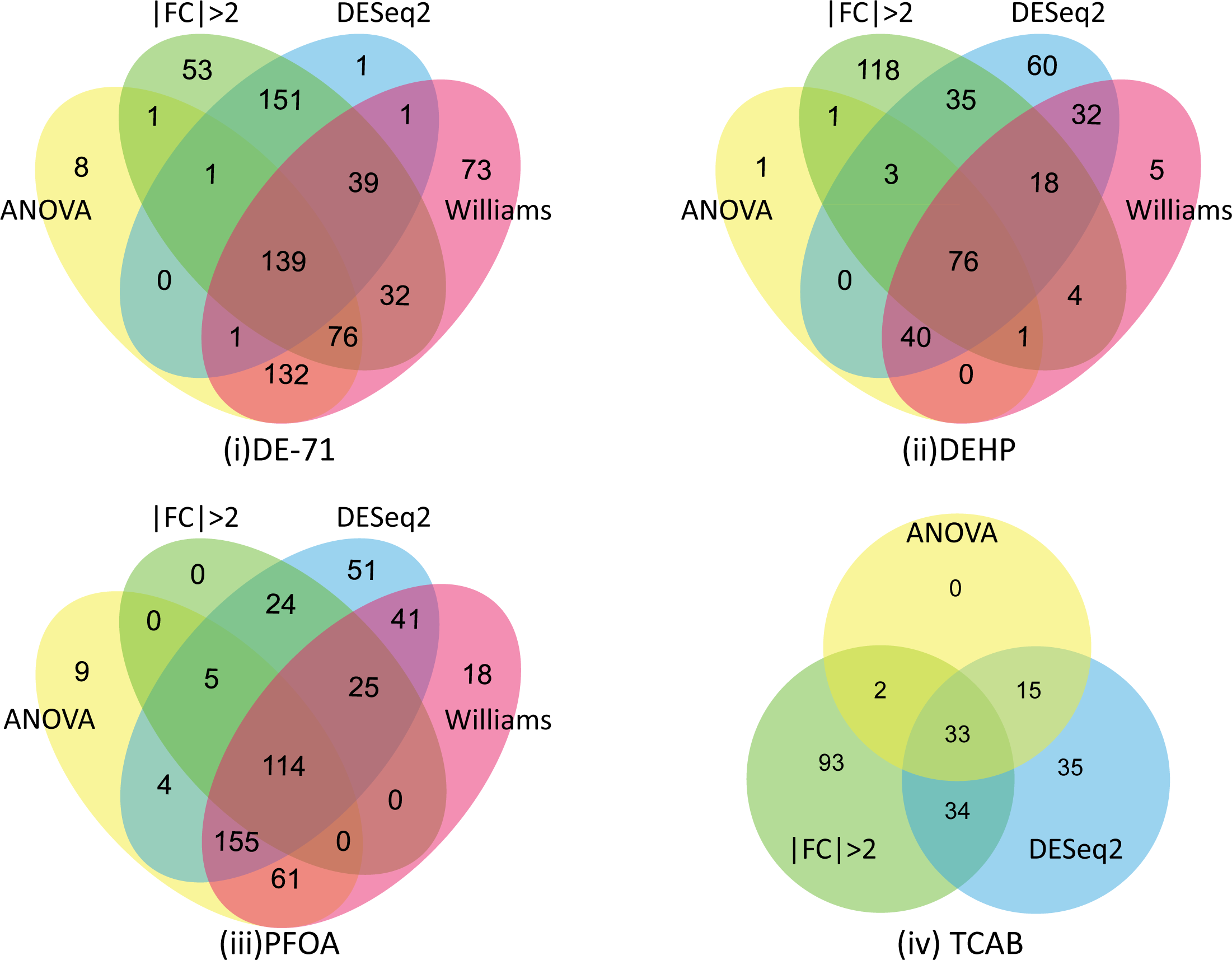
Distribution of significant genes (FDR<0.05 & |FC|>1.5) which also had a BMD best fitting model that passed model QC assessment (BMD <maximum experimental dose & goodness of fit p-value <0.1 & BMDU/BMDL <40) from in vivo rat liver samples. The Venn diagrams for TCAB do not include any genes for the WTT as no genes met these criteria. Acrylamide and BDCA are not shown as they had no significant genes by ANOVA nor by Williams Trend Test.

**Supplemental figure 3.**
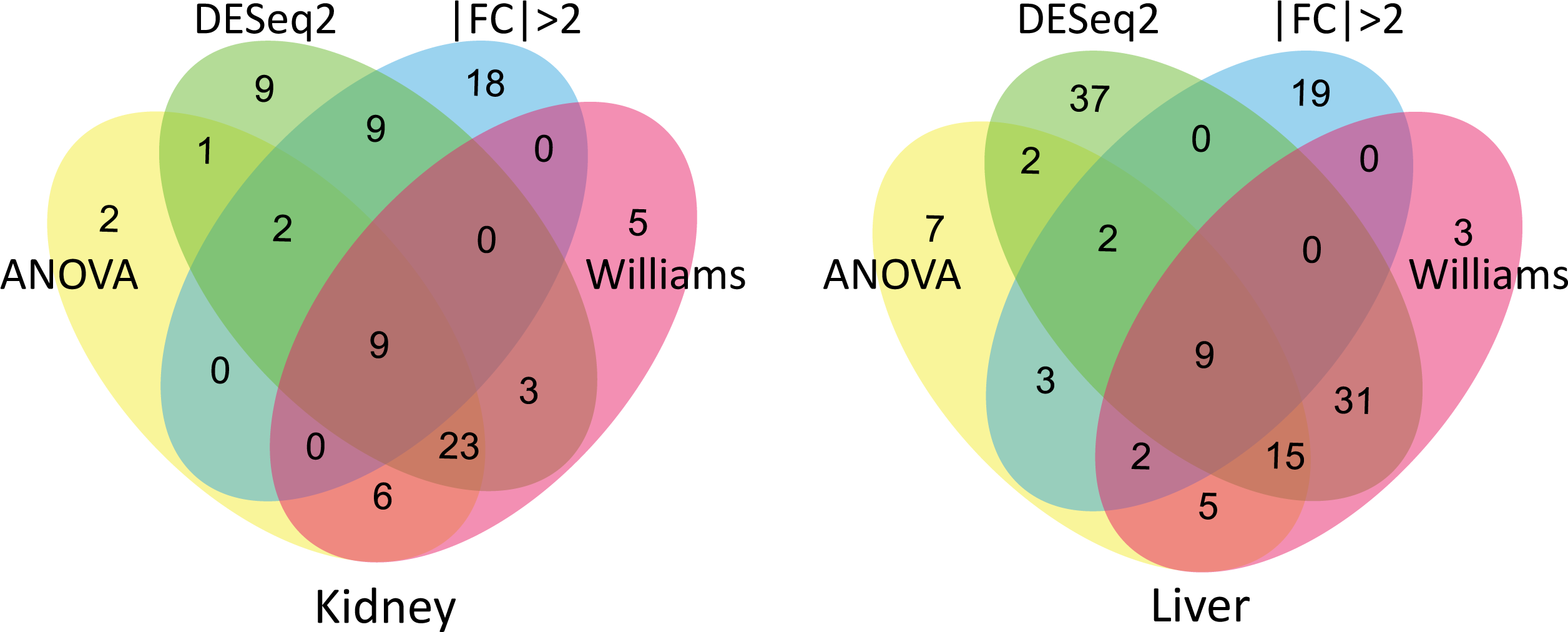
PFOA: Distribution of enriched pathways from BMD gene enrichment analysis. A pathway was considered enriched if it at least 5 query genes were found amongst the pathway elements, and the over-representation analysis Fisher’s exact p-value < 0.05. DESeq2 detected many more unique enriched pathways than any other metric. Reactome is hierarchical and thus cellular pathways progress from very general parent categories (e.g. *Metabolism*) to ever more refined child processes, terminating with discrete functional processes such as *Cholesteral biosynthesis*. PFOA allowed for the most robust comparison across the significantly differentially expressed gene sets as with all gene sets there were more than 20 enriched pathways for both liver and kidney samples. Given the differences in the distribution of significantly expressed genes detected by each metric, each also results in some enriched pathways unique to the definition of significance for differential gene expression. In the livers of animals treated with PFOA, genes identified using ANOVA, WTT, and DESeq2 produced significantly enriched pathways in the same primary functional categories in Reactome (*Cell Cycle, Metabolism, Signal transduction, Immune System, Gene expression, Programmed cell death, Cellular response to external stimuli, Extracellular matrix organization, and Developmental biology*) but with differing numbers of child processes detected within these parental processes. Pathways enriched in the kidney also included many of the same primary pathway categories in Reactome as seen with liver samples, and again the primary difference between the four differential expression metrics was in the numbers of child processes detected as enriched within those large parental Reactome categories. Since the likelihood of detecting significantly enriched processes is in part a function of the gene lists available for input into an enrichment analyses, the sensitivity of a differential gene expression metric has implications for the sensitivity of the enrichment analysis.

**Supplemental figure 4.**
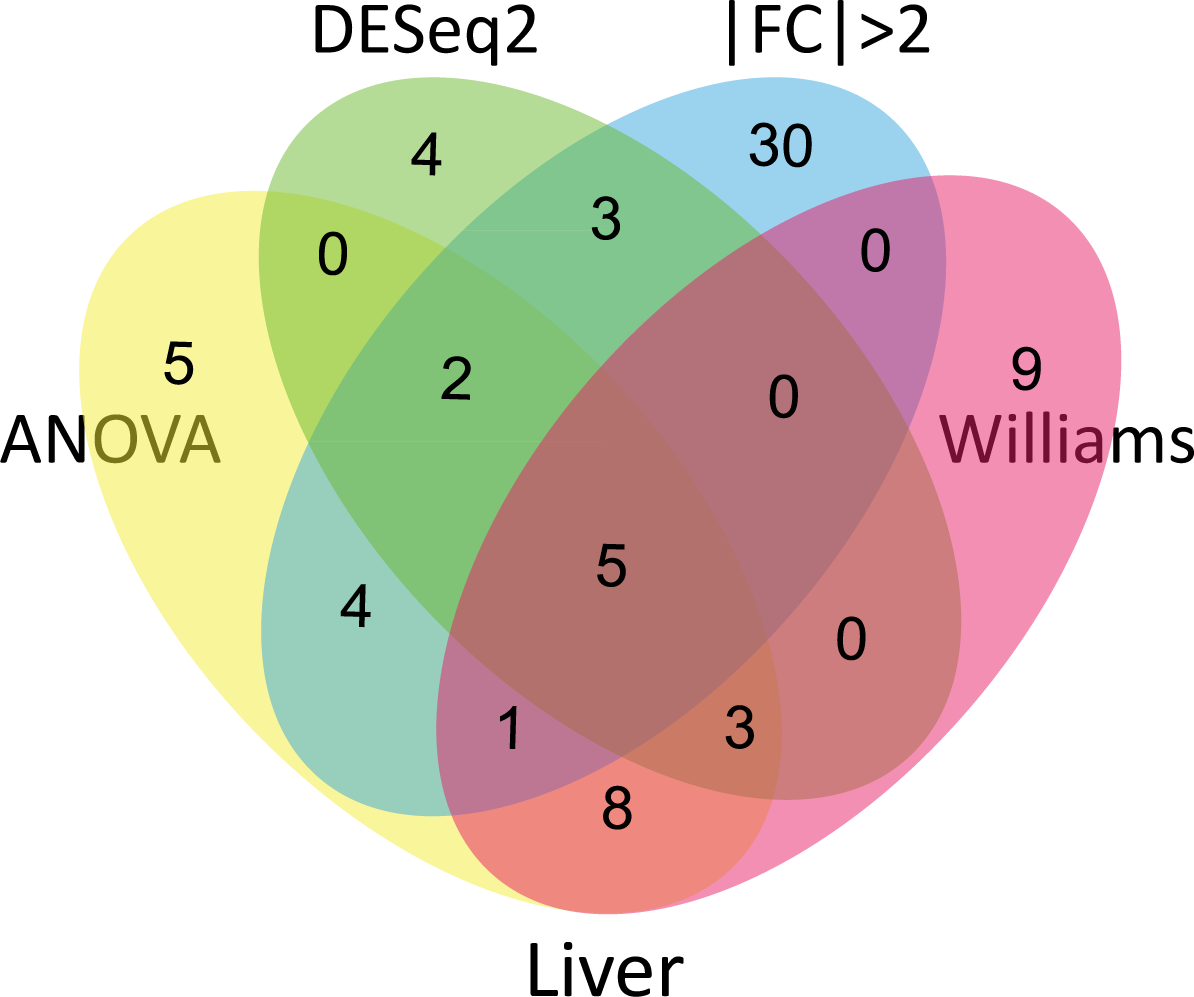
DE-71: Distribution of enriched pathways from BMD gene enrichment analysis. A pathway was considered enriched if it at least 5 query genes were found amongst the pathway elements, and the over-representation analysis Fisher’s exact p-value < 0.05. Genes significant by fold change only include many more pathways than enriched with any of the statistical metrics, but have the least overlap with pathways enriched using genes significant by any other metric.

**Supplemental figure 5.**
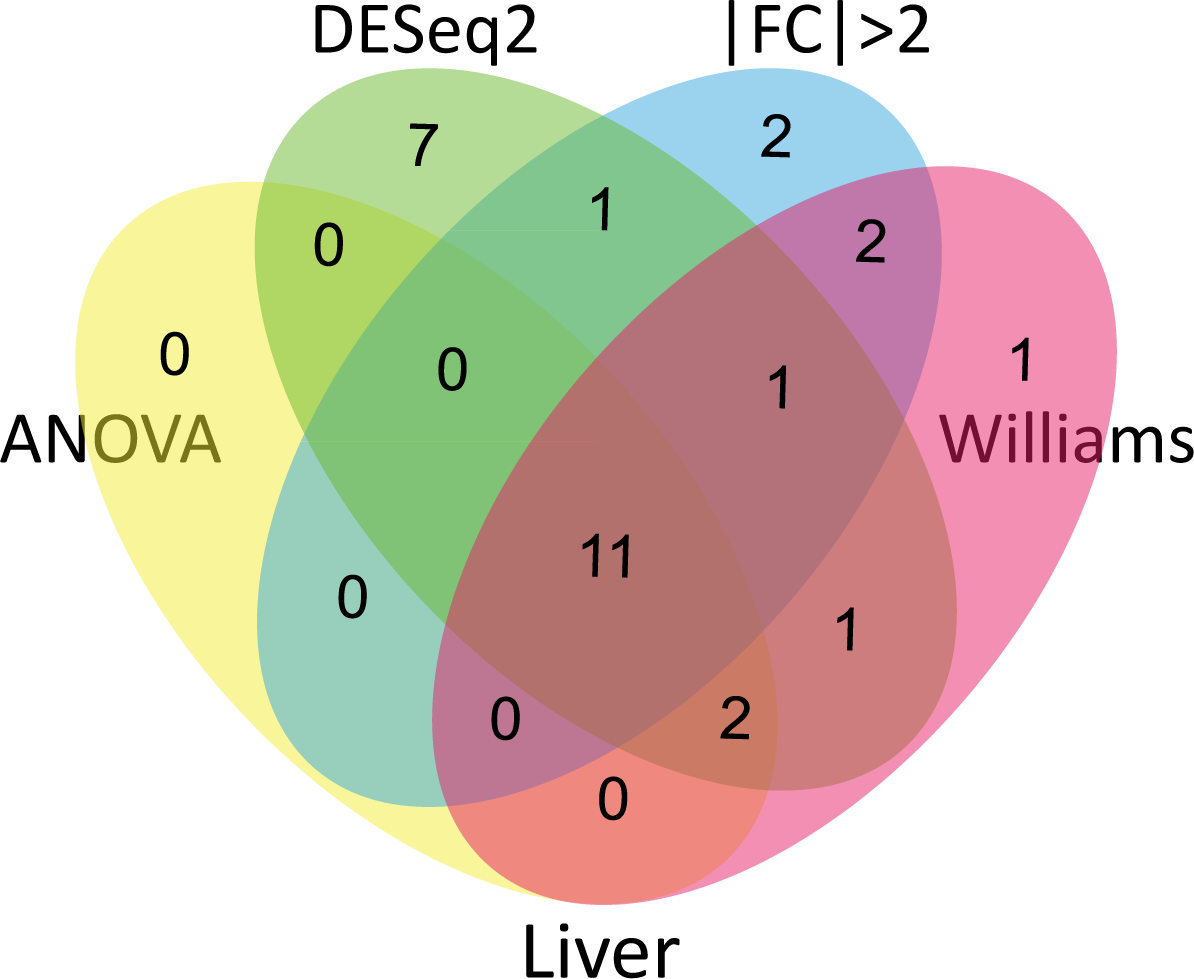
DEHP in liver: Distribution of enriched pathways from BMD gene enrichment analysis. A pathway was considered enriched if it at least 5 query genes were found amongst the pathway elements, and the over-representation analysis Fisher’s exact p-value < 0.05. This analyses indicated the greatest overlap in pathways amongst all four gene significance metrics (11 pathways enriched in all four analyses), but genes significant by DESeq2 also resulted in 7 pathways not enriched with genes from any of the other three metrics.

